# Formation of the clamped state by Scc2 and DNA overcomes the constraints imposed by zipping-up of the SMC coiled coils on cohesin’s ATPase

**DOI:** 10.1101/2022.06.19.496727

**Authors:** Menelaos Voulgaris, Kim A Nasmyth, Madhusudhan Srinivasan

## Abstract

In addition to mediating sister chromatid cohesion, cohesin, by virtue of Loop Extrusion (LE), organises the spatial arrangement of interphase DNA. The latter activity relies on DNA and Scc2 dependent ATP hydrolysis by cohesin. How the impetus from the ATPase cycle translates into reeling of DNA loops into the SMC kleisin rings is still unclear. The SMC coiled coils show several striking structural features like folding and zipping-up, if and how these structural states affect cohesin’s activity is still unclear. We show here that cohesin’s loop extruding motor contains an internal constraint that regulates its ATPase activity, zipping-up of the coiled coils impedes ATP hydrolysis by cohesin. We show that integrity of a region where the coiled coils emerge for the SMC hinge domains, SMC ‘wrist’, is critical for the zipping up of the coiled coils and the resulting inhibition of cohesin’s ATPase. Clamping of DNA by Scc2 onto the engaged SMC heads in the presence of ATP leads to unzipping of the coiled coils and permits ATP hydrolysis. Strikingly, irreversible folding of the coiled coils at the elbow region does not lead to any measurable change to the ATPase activity suggesting that recurrent cycles of folding and unfolding of the coiled coils is not necessary for driving continuous ATP hydrolysis by cohesin.

## Introduction

Individualisation of replicated DNAs is critical to ensure accurate segregation of chromosomal DNAs, which is fundamental for cell proliferation. In most if not all life forms, this monumental feat is accomplished by a set of highly conserved molecular machines called the SMC-Kleisin protein complexes. SMC complexes are topological devices and their ability to organise chromosomal DNAs is thought to stem from their ability to perform Loop Extrusion (LE), an activity that involves processive enlargement of DNA loops in an ATP hydrolysis dependent manner (Davidson and Peters, 2021; Yatskevich et al., 2019). Because of temporal separation of S phase from mitosis eukaryotic cells face an additional problem. Tethering of individualised sister chromatids from S phase until their disjunction during mitosis is critical for eukaryotic chromosome segregation, this is achieved by an eukaryote specific SMC complex called cohesin, which in addition to organising chromosomal DNAs by LE (Davidson et al., 2019; Kim et al., 2019), confers sister chromatid cohesion by co-entrapment of sister DNAs within the SMC-Kleisin ring (Guacci et al., 1997; Michaelis et al., 1997; Srinivasan et al., 2018).

The core of SMC complexes is a tripartite ring composed of two SMC proteins and an α-kleisin subunit (Haering and Gruber, 2016), with HEAT repeat containing regulatory subunits associating with the kleisin (Wells et al., 2017). While the geometry of all SMC complexes is remarkably similar, since this study is based on analysis of cohesin, we will henceforth refer to and elaborate on the structural features of cohesin. The cohesin ring is composed of Smc1 and Smc3 proteins associated with the kleisin Scc1(Gruber et al., 2003). The Smc1 and Smc3 proteins have two globular domains, the hinge and ATPase head domains, separated by long coiled coils. The ABC-like ATPase head domain contains a Walker A motif that binds ATP, a signature motif capable of binding the γ-phosphate of ATP bound to an adjacent ATPase head as well as a Walker B motif necessary for ATP hydrolysis (Gligoris et al., 2014; Haering et al., 2002). As in the case of proteins with ABC-like cassettes, cohesin’s ATPase activity arises from sandwiching of two ATP molecules between engaged ATPase heads (E heads) prior to their hydrolysis (Lammens et al., 2004). When cohesin’s ATPase heads are disengaged, the coiled coils of Smc1 and Smc3 associate with each other along much of their length (Petela et al., 2021). When this ‘zipping up’ includes the sections of coiled coils close to the ATPase heads, it forces them to adopt a configuration in which they are juxtaposed, a state that is distinct from, and incompatible with ATP dependent head engagement (J Heads) (Chapard et al., 2019). In addition to their tendency to zip up, the coiled coils fold around the ‘elbow’ (Burmann et al., 2019); a discontinuity in the coiled coils around which the SMC proteins fold (Figure 1A), this results in the association of the hinge domain to a region of the coiled coil that is in proximity to the ‘joint’; another discontinuity in the coils close to the ATPase heads (Figure 1A) (Petela et al., 2021).

**FIGURE 1.**
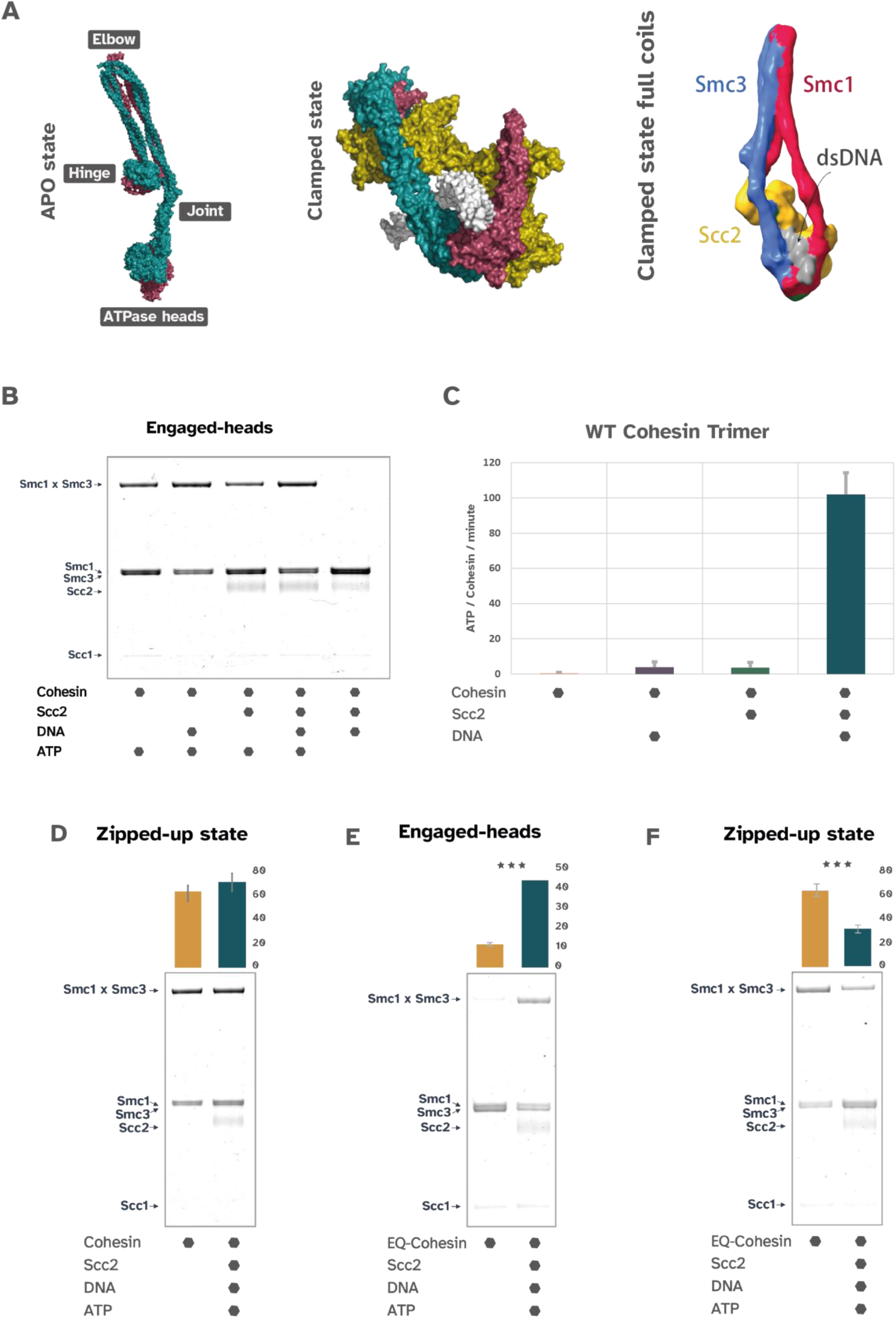
A. Cryo-EM structures of the APO (7OGT *I* personal communication) and clamped state (6ZZ6) of cohesin. Clamped state with full coiled coils as seen in Collier et al, 2020. B. In vitro cystein crosslinking of Smc1 N1192C and Smc3 R1222C measuring the Smc head association indicative of the engeged heads confromation. C. WT cohesin trimer ATPase activity measurements. ATP hydrolysis by the cohesin complex was measured in 4 different conditions. D. In vitro cystein crosslinking to measure the formation of the Zipped up state of WT cohesin trimer in the presence or absence of Scc2, DNA and ATP. The experiment was repeated 3 times and the quantification is shown in the bar graphs above the gels (p<0.001). E. In vitro cystein crosslinking to measure the formation of the Engaged state of EQ cohesin trimer in the presence or absence of Scc2, DNA and ATP. The experiment was repeated 3 times and the quantification is shown in the bar graphs above the gels (p<0.001). F. In vitro cystein crosslinking to measure the formation of the Zipped up state of EQ cohesin trimer in the presence or absence of Scc2, DNA and ATP. The experiment was repeated 3 times and the quantification is shown in the bar graphs above the gels (p<0.001).

Cohesin’s abilities to associate with DNA, to extrude DNA loops and to tether sister DNAs is regulated by a group of proteins with remarkably similar structure called the HAWKS. Cohesin’s HAWKS are Scc3, Scc2 and Pds5. While Scc3 is associated constitutively, Scc2 and Pds5 are mutually exclusive. Pds5 is essential for stable maintenance of sister chromatid cohesion in post replicative cells (Hartman et al., 2000; Panizza et al., 2000). Scc2 is essential for loading and maintenance of cohesin’s association with un-replicated DNA but is dispensable for maintaining pre-established cohesion in post replicative cells (Ciosk et al., 2000; Srinivasan et al., 2019). In vitro, ATP hydrolysis by cohesin is strictly dependent on Scc2 (Petela et al., 2018). Scc2 is also necessary for continuous LE by cohesin in vitro, presumably to catalyse continuous ATP hydrolysis by cohesin (Davidson et al., 2019). While sister chromatid cohesion and LE might be separable activities of cohesin, the fact that Scc2 is essential for topological as well as non-topological association of cohesin with DNA suggests that the two activities of cohesin must share common intermediates (Collier et al., 2020; Srinivasan et al., 2018). Recent cryo-EM studies suggest that one such intermediate is a ‘clamped state’ of cohesin (Figure 1A), where DNA is clamped onto the ATP bound SMC head domains by Scc2 (Collier et al., 2020; Higashi et al., 2020; Shi et al., 2020). Crucially, formation of the clamped state in vitro does not require Scc3, which is essential for cohesin’s stable association and entrapment of DNA in vitro and in vivo, as well as for LE by cohesin in vitro (Collier et al., 2020; Davidson et al., 2019; Losada et al., 2000; Roig et al., 2014). Cohesin that is defective in ATP hydrolysis due to mutations in the Walker B motif of the SMC ATPase domains is incapable of stable association or entrapment of chromosomal DNA but accumulates at the centromeric loading sites on yeast chromosomes (Hu et al., 2011; Srinivasan et al., 2018). In vitro, the walker B mutant cohesin forms and accumulates in the clamped state (Collier et al., 2020; Higashi et al., 2020; Shi et al., 2020). The fact that the clamped state is detected in other SMC complexes suggests that clamping of DNA is a key intermediate step in the enzymatic activity of all SMC complexes (Burmann et al., 2021; Lee et al., 2022).

While the molecular mechanism of LE is still unclear, all models for LE envision recurrent formation and dissolution of the clamped state (Bauer et al., 2021; Davidson and Peters, 2021; Hassler et al., 2018; Higashi et al., 2021; Yatskevich et al., 2019). Single molecule FRET data suggests that the formation and dissolution of the clamped state are coordinated with substantial conformation changes to the cohesin ring involving unzipping and unbending of the colied coils (Bauer et al., 2021). Whether and how these conformation changes affect cohesin’s ATPase activity and LE is still unclear. In the absence of ATP, cohesin is in a ‘zipped up’ and folded conformation (Figure 1A). Under this condition Scc2 remains associated with the disengaged ATPase heads and the N-terminus of Scc2 interacts with the SMC hinge domain (Petela et al., 2021). In the clamped state, the coiled coils, at least in a region proximal to the ATPase heads is unzipped (Collier et al., 2020; Higashi et al., 2020; Shi et al., 2020) (Figure 1A). Biochemical analysis suggest that the cohesin can attain the ATP dependent head engaged state, at least transiently, in the absence of both Scc2 and DNA and cryo-EM studies of a truncated cohesin complex shows that the ATP bound head domains can engage in the absence of DNA and Scc2, with the coiled coils in an unzipped state (Bauer et al., 2021; Collier et al., 2020; Muir et al., 2020). This raises the question as to whether ATP dependent head engagement per se, without the formation of the clamped state is sufficient for the coiled coil unzipping. However, both Scc2 and DNA are critical for cohesin’s ATPase activity in vitro (Petela et al., 2018). Another feature of cohesin’s coiled coils that has emerged from the structural studies is its folding at the elbow. Cohesin’s coiled coils remain folded in the clamped state, suggesting that clamping and folding of the coiled coils cannot be mutually exclusive states (Collier et al., 2020; Shi et al., 2020). Crucially, the effect of the coiled coil conformations, namely their zipping up and folding on cohesin’s ATPase activity is still unclear. Is there a relationship between the coiled coil conformation and cohesin’s ATPase cycle? What is the role of Scc2 and DNA in promoting ATP hydrolysis and why can’t cohesin hydrolyse ATP in their absence are key questions. Given that LE depends on continuous ATP hydrolysis by cohesin, answering these questions is critical to begin understanding the molecular mechanism of LE. In the present study we address the regulation of cohesin’s ATPase activity and probe the relationship between the conformation changes to cohesin ring with its ATPase cycle. We show that the zipping up of the coiled coils impedes ATP hydrolysis by cohesin in the absence of DNA and Scc2. ATP-dependent engagement of cohesin’s Smc1 and Smc3 head domains and clamping of DNA by Scc2 onto the engaged heads leads to unzipping of the coiled coils and permits ATP hydrolysis. We identify a section of the coiled coils that are critical for imposing the constraint on cohesin’s ATPase; cohesin ‘wrist’. Compromising the integrity of cohesin wrist emancipates cohesin’s ATPase from Scc2 and DNA, but results in cell inviability. This suggests that the internal constraint present within cohesin’s loop extruding motor ensures that ATP hydrolysis happens only under the right conditions. Incredibly, while zipping up of the coiled coils in incompatible with ATP hydrolysis, we find that cohesin is fully functional as an ATPase in the folded confirmation. This implies that recurrent rounds of folding and unfolding are not required for continuous ATP hydrolysis by cohesin.

## Results and Discussion

A major difference between the cohesin in its ‘apo’ state bound to Scc2 in the absence of ATP and the ATP-bound clamped state is the conformation of the Smc coiled coils. Though the coiled coils are folded in both cases, they are zipped up in the absence of ATP and DNA but held agape at least up to the joint when DNA is clamped by Scc2 onto the ATPase heads (Figure 1A). Unzipping of the coiled coils could be driven by head engagement per se. Alternatively, it might additionally require the clamping of DNA to engaged heads by Scc2. To understand the effect of colied coil conformation, Scc2 and DNA in regulating ATP hydrolysis by cohesin, we use BMOE dependent crosslinking of carefully positioned cysteine residues as a reporter of the conformation of the ATPase heads upon engagement, juxtaposition (Chapard et al., 2019), coiled coil folding (Petela et al., 2021) and zipping-up (Chapard et al., 2019) (Figure S1A). The cysteine residues have all been validated in vivo, the mutations do not affect cell viability and do not hinder the normal function of cohesin (Chapard et al., 2019; Petela et al., 2021). We expressed and purified the wild type and cysteine substituted yeast cohesin trimers (Smc1, Smc3, Scc1) along with Scc2 using the baculovirus expression system in SF9 insect cells (Figure S1B, see methods for purification conditions).

### Formation of the clamped state leads to unzipping of the coiled coils emanating from the ATPase heads

While cohesin’s ATPase heads can engage in the presence of ATP alone (Figure 1B), ATP hydrolysis by cohesin is strictly dependent upon the presence of both DNA and Scc2 (Figure 1C) (Petela et al., 2018). In the absence of ATP, cohesin’s coiled coils are zipped-up along their entire length (Figure 1A) (Petela et al., 2021). To measure zipping-up of the coiled coils, we treated the cohesin trimer with cysteine residues in the coiled coil near the joint region with BMOE. This revealed that zipping-up, as measured by crosslinking of the coiled coils was always detected and addition of ATP, DNA and Scc2 did not reduce the crosslinking efficiency (Figure 1D). This could be because under conditions cohesin can hydrolyse ATP, the clamped state (where the coiled coils are unzipped) is too transient in the ATPase cycle of cohesin to enable measurement of the change of state by BMOE crosslinking. If this is the case, the walker B mutant EQEQ cohesin that accumulates in the clamped state should permit measurement of the effect of clamping on coiled coil conformation. To do this, we purified EQEQ cohesin containing the head engagement and coiled coil zipping-up reporter cysteine substitutions (Figure S1B). We noticed that unlike the wild type cohesin, where it depends only on ATP, head engagement as measured by BMOE crosslinking of the EQEQ cohesin depends on the presence of ATP, Scc2 and DNA (Figure 1E). We next addressed the effect of clamping on coiled coil zipping-up using the EQEQ cohesin. Interestingly, we measured a significant reduction of the zipping-up of the coiled coils up to the joint region only in the presence of ATP, DNA and Scc2 (Figure 1F). While ATP bound wild type cohesin head domains attain the engaged state without DNA and Scc2 (Figure 1B), this does not lead to ATP hydrolysis or a measurable change in the coiled coil zipping-up (Figure 1C and 1D). However, the EQEQ cohesin that accumulates in the clamped state in the presence of ATP, DNA and Scc2 shows a marked reduction in zipping-up of the coiled coils at the joint. This suggests stable clamping of DNA by Scc2 on ATP bound heads leads to unzipping of the coiled coils, at least up to the joint. Clamping and zipping-up of the coiled coils are therefore likely to be mutually exclusive states. We therefore asked whether unzipping of the coiled coils is necessary for cohesin’s ATPase activity.

### Zipping-up of the coiled coils at the joint region impedes cohesin’s ATPase activity

If unzipping of the coiled coils is necessary for cohesin’s ATPase activity, cohesin that is trapped in a state where the coiled coils beneath the joint are irreversibly zipped-up should be incapable of ATP hydrolysis. To test this, we reasoned that we could measure the ATP hydrolysis of pre-crosslinked cohesin containing the cysteine substitutions that report coiled coil zipping-up. This reasoning assumes that BMOE treatment per se, does not adversely affect cohesin’s ATPase activity. To check if this is the case, we pre-treated wild type cohesin that does not contain any cysteine substitutions with BMOE. Crucially, this treatment does not result in any nonspecific protein crosslinks of the wild type cohesin (Figure 2A). We measured the rate of ATP hydrolysis by the wild type cohesin that was pre-treated with either BMOE or a DMSO control, which revealed that cohesin was fully proficient in ATP hydrolysis under both conditions (Figure 2A). This strongly suggests that BMOE treatment per se does not adversely affect cohesin’s ATPase activity. Cohesin, when not bound to ATP is in a state where the ATPase heads are juxtaposed (Petela et al., 2021), this J-state is mutually exclusive with the head engagement (E-state) (Chapard et al., 2019) and therefore cohesin trapped in the J state should be compromised for ATP hydrolysis. We found that this is indeed the case, treatment of cohesin containing J-state reporter cysteines with BMOE results in about 70% of the cohesin molecules becoming irreversibly stuck in the J-state, and a corresponding decrease in ATP hydrolysis by the pre-crosslinked cohesin (Figure 2A and 2B). Interestingly, we found that the cohesin complexes that were irreversibly stuck in the zipped-up state at the joint region behaved very similar to the J-state cohesin and were incapable of ATP hydrolysis, treatment with BMOE resulted in about 68% of cohesin molecules being stuck in the zipped-up state, resulting in a corresponding decrease in the rate of ATP hydrolysis by pre-crosslinked cohesin (Figure 2A and 2B). This observation implies that when coiled coils are zipped-up at the joint region, cohesin is incapable of ATP hydrolysis. While ATP is sufficient to induce head engagement, both Scc2 and DNA are required for attaining the clamped state, unzipping of the coiled coils up to the joint, and ATP hydrolysis

**FIGURE 2.**
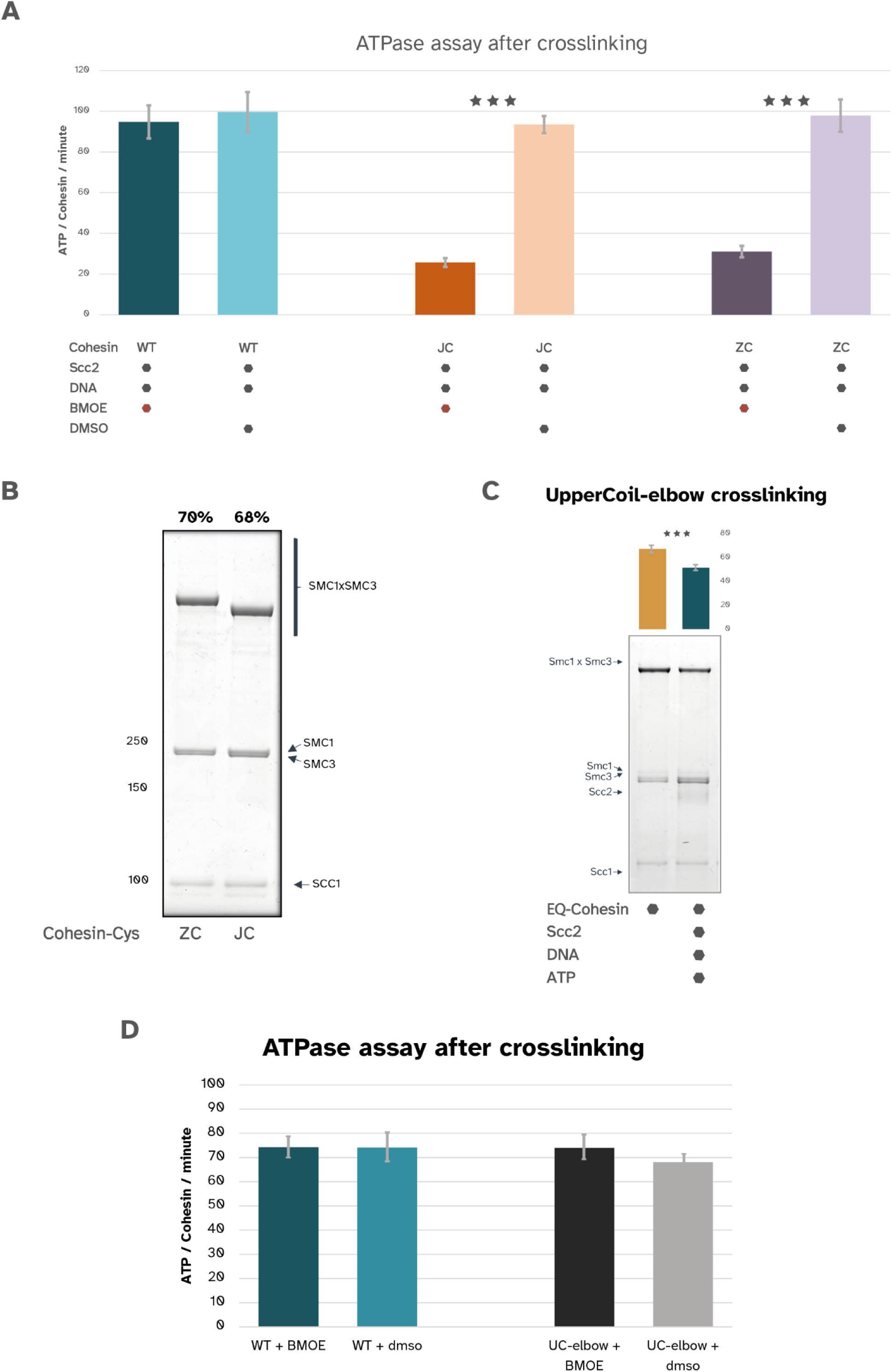
A. ATPase activity measurements after cmsslinking with BMOE in comparison with DMSO controls. Cohesin trimers with cysteine pairs capturing the J-head state (JC) and the zipped-up confmmation (ZC) were tested along with (WT). Each experiment was repeated 3 times (p<0.001). B. In vitro cystein cmsslinking to measure the efficiency of the cmsslinking either in the Zipped up state (ZC) or the J-head state (JC) before release in the ATPase assay. C. In vitm cystein cmsslinking to measure the formation of the Upper-Coil zipping up of EQ cohesin trimer in the presence or absence of Scc2, DNA and ATP. The experiment was repeated 3 times and the quantification is shown in the bar graphs above the gels (p<0.001). D. ATPase activity measurements after cmsslinking with BMOE in comparison with DMSO controls. Cohesin trimers with cysteine pairs capturing the Upper-coil zipped up state (UC-elbow) and were tested along with (WT). Each experiment was repeated 3 times (p<0.001).

In the absence of ATP, the coiled coils are zipped-up along their entire length. Unzipping of the coils up to the joint region is necessary for ATP hydrolysis. How far along the length does this unzipping have to happen? To address this, we engineered cysteine pairs at the elbow region (Figure S1A and S1B) to trap the coils in a zipped-up state at the elbow. Treatment of EQEQ cohesin containing the cysteine substitutions at the elbow region resulted in crosslinking of about 60% of the cohesin molecules, addition of Scc2, DNA, and ATP resulted only in a modest decrease in crosslinking (Figure 2C). Crucially, more than 50% of the molecules remain in the zipped-up state under this condition (Figure 2C). We asked if crosslinking the coiled coils at the elbow region affects the ATPase activity. To do this, we measured ATP hydrolysis by pre-crosslinked complexes, this revealed that crosslinking of coiled coils at the elbow did not affect the rate of ATP hydrolysis, cohesin molecules irreversibly crosslinked at the elbow were fully competent in hydrolysing ATP (Figure 2D). This implies that while unzipping of the coiled coils at the joint region is crucial for it, unzipping at the elbow region is not required for Scc2 and DNA mediated stimulation of ATP hydrolysis. In other words, the zipping up of the coiled coils around the joint imposes a constraint on the ATPase domain that must be overcome by Scc2 and DNA to permit ATP hydrolysis.

### Permanent unzipping of the coiled coils permits DNA and Scc2 independent ATP hydrolysis by cohesin

In the absence of ATP, the coiled coils are zipped up along their entire length (Figure 1A) (Petela et al., 2021). What features determine this alignment of cohesin’s coil? Since the coils at the elbow region are rarely unzipped, and the fact that irreversible zipping at the elbow does not inhibit cohesin’s ATPase activity (Figures 2C and 2D), we asked whether an engineered cohesin with coiled coils extending up to the elbow region would be able to adopt a rigid enough zipped-up state at the joint region, capable of inhibiting ATP hydrolysis. We therefore expressed and purified a truncated cohesin trimer, where both SMCs extend up to the elbow region with their N terminal ascending and the C terminal descending coils connected with a flexible GS linker (Figure S1B). The truncated cohesin complex, was first assessed for its ability to crosslink in the zipped-up state at the joint region using the cysteine reporters. Treatment of the full length cohesin with BMOE in the absence of ATP leads to crosslinking of the coils at the joint region in about 60% of molecules (Figure 3A). In contrast, the same treatment of the truncated complex did not yield almost any measurable crosslinking (Figure 3A). Interestingly, unlike the full length cohesin (shown as narrow, lightly shaded bars) where the ATPase activity is strictly dependent on both Scc2 and DNA, we found that truncated cohesin was fully active as an ATPase, even in the absence of DNA (Figure 3B). Even more strikingly, the truncated complex retained significant ATPase activity in the absence of Scc2 (Figure 3B). Remarkably, removing the constrains imposed by the zipped-up coiled coils on the ATPase domain abolishes the strict requirement of DNA and to a large extent, Scc2 for ATP hydrolysis.

**FIGURE 3.**
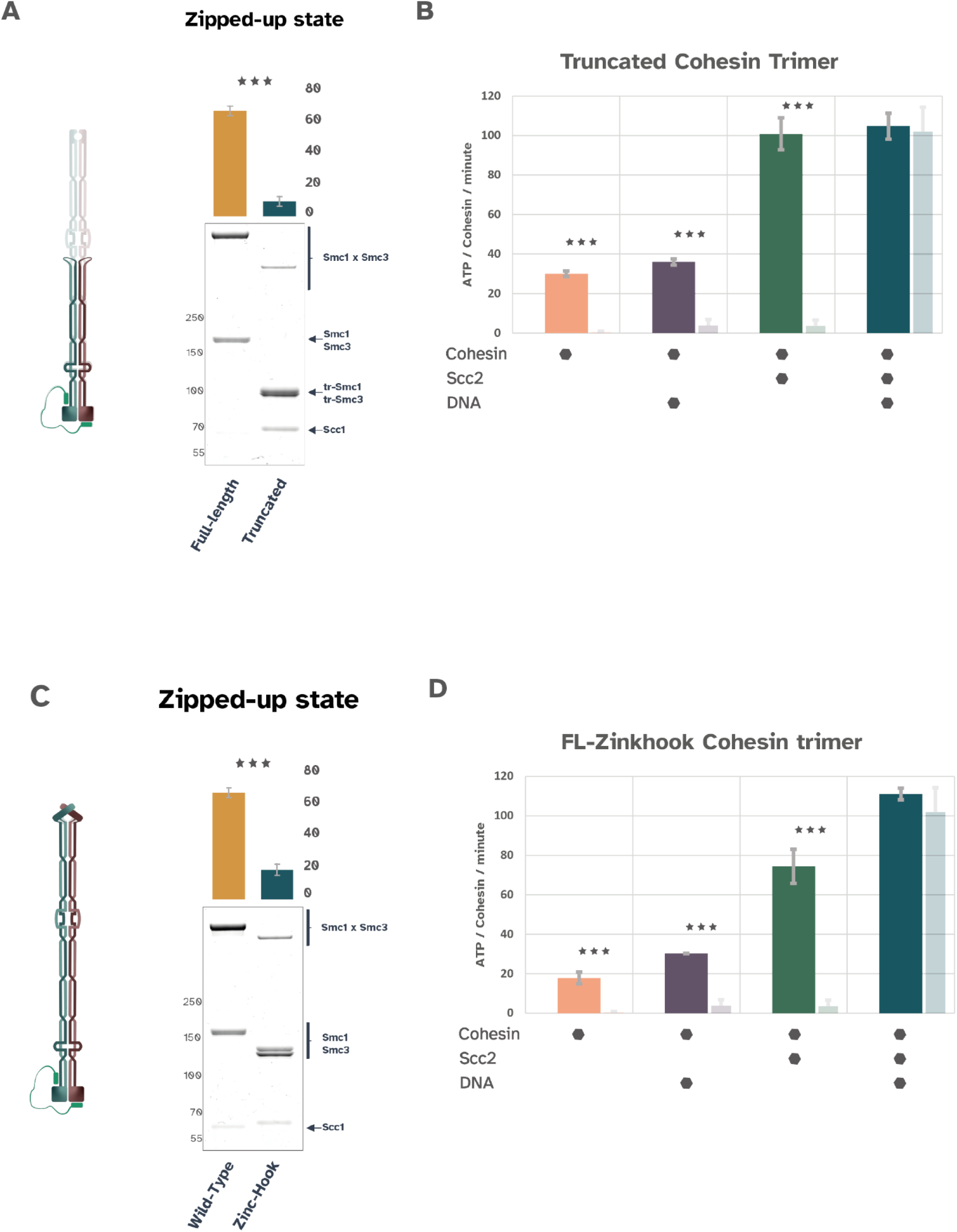
A. In vitro cystein crosslinking to measure the formation of the Zipped up state of *WT* full length cohesin trimer versus the truncated trimer. The experiment was repeated 3 times and the quantification is shown in the bar graphs above the gels (p<0.001). B. Truncated cohesin trimer ATPase adivity measurements. ATP hydrolysis by the truncated cohesin complex was measured in 4 different conditions. Light shade bars represent *WT* full length cohesin’s ATPase adivity in each condition. C. In vitro cystein crosslinking to measure the formation of the Zipped up state of *WT* full length cohesin trimer versus the “Zink-hook” trimer. The experiment was repeated 3 times and the quantification is shown in the bar graphs above the gels (p<0.001). D. “Zink-hook” cohesin trimer ATPase adivity measurements. ATP hydrolysis by the “zink-hook” cohesin complex was measured in 4 different conditions. Light shade bars represent *WT* full length cohesin’s ATPase adivity in each condition.

Clearly, the coiled coils up to the elbow region are incapable of adopting the zipped-up conformation. Zipping-up could therefore require the full-length coiled coils and/or the hinge domain. We were unsuccessful in purifying cohesin lacking the hinge but containing the full-length coils. However, we could purify a form of cohesin that contained the entire coiled coil domain but was held together by the Rad50 zinc hook (ref) instead of the hinge domain. We next asked if the full-length coiled coils are sufficient for cohesin adopting the zipped-up conformation. We found that the Rad50 zinc-hook containing cohesin (Figure S1B) was significantly compromised in coiled coil zipping-up in the absence of ATP (Figure 3C) suggesting that the full-length coiled coil domain is not sufficient to induce coiled coil zipping in the absence of ATP. Interestingly, we found that, like thr truncated elbow length cohesin, the Rad50 zinc-hook containing cohesin was able to hydrolyse ATP without DNA, and retained significant activity in the absence of Scc2 (Figure 3D). One can derive two interesting implications from the results described above: 1. Cohesin’s hinge domain is critical for the zipping-up of coiled coils at the joint region. 2. Compromising the ability of the coiled coils to zip-up at the joint results in relaxation of the strict requirements of Scc2 and DNA for ATP hydrolysis. This is consistent with the notion that Scc2 by clamping DNA onto the ATPase heads leads to unzipping of the coils up to the joint and promotes ATP hydrolysis by cohesin.

### The transition region between cohesin’s hinge and coiled coils: ‘cohesin wrist’ region is critical for coiled coil zipping up

Results described above suggest a role for the hinge domain in enabling the zipped-up conformation of the coiled coils. Cryo-EM structure of cohesin in the absence of ATP shows that the hinge domain adopts a conformation where it is almost at a right angle with the emerging coiled coils (Figure 4A) (Petela et al., 2021). In this conformation, the region where the hinge domain transitions into the coiled coils associates with the zipped-up coiled coils near the joint (Figure 4A) (Petela et al., 2021). We therefore asked whether the hinge domain per se, or the specific way the hinge domain transitions into the coiled coils was responsible for the zipping-up. To this end, we introduced a flexible glycine-serine linker between the hinge and the coiled coils. We noticed that this led to a significant reduction in the zipping-up of the coils at the joint (Figure 4B) and DNA independent ATP hydrolysis, with significant ATP hydrolysis activity in the absence of Scc2 (Figure 4C), suggesting that it is not the hinge domain per se that is important for the coiled coil zipping-up. Instead, the integrity of the region where the hinge domain transitions into the coiled coil is crucial for zipping-up of the coils and the resultant constraint on the ATPase domain.

**FIGURE 4.**
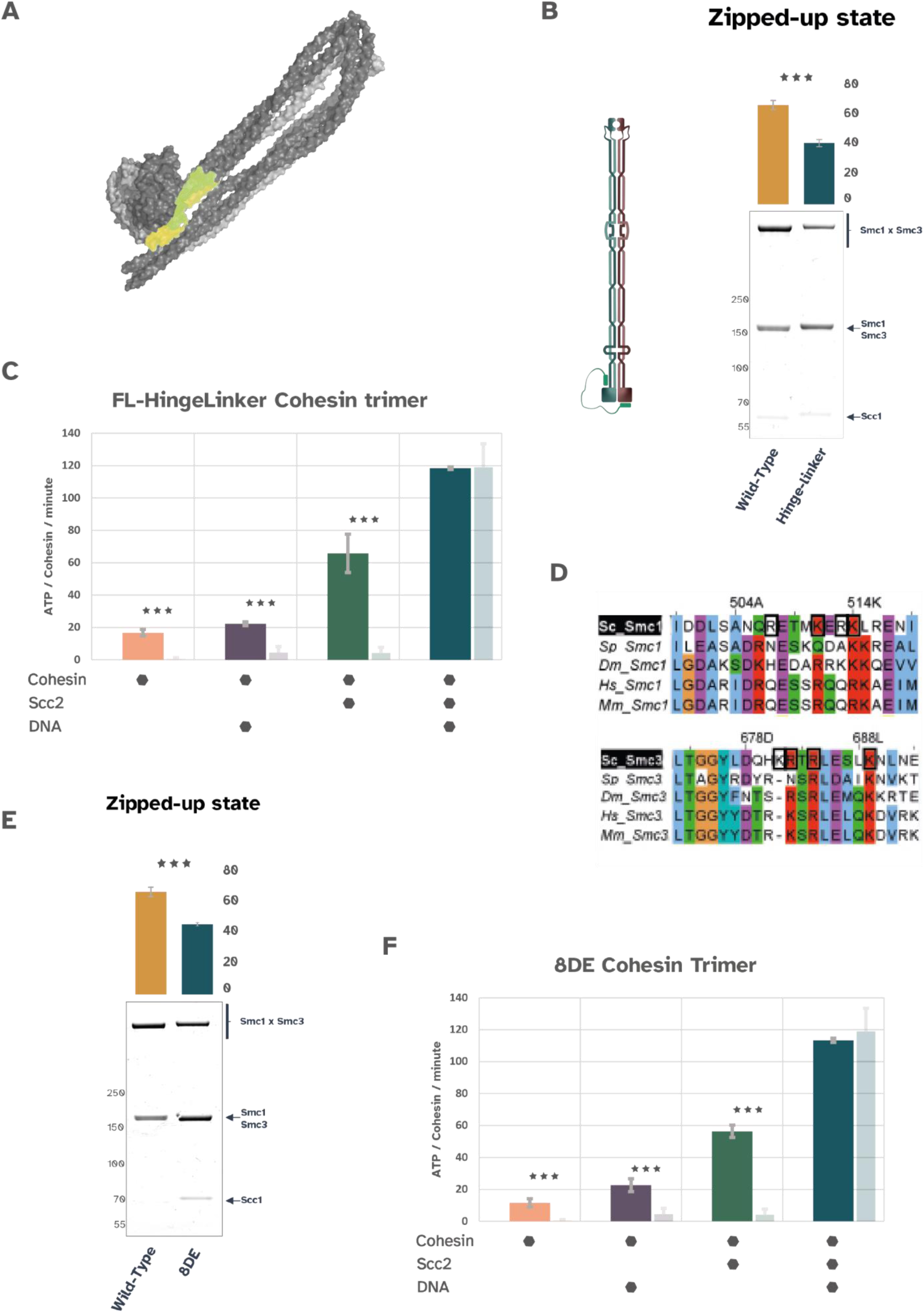
A. Cryo-EM structures of the cohesin coiled-coils/ hinge in the folded conformation. The cohesin wrist is indicated in green hue. B. In vitro cystein crosslinking to measure the formation of the Zipped up state of WT full length cohesin trimer versus the “Hinge-linker” trimer. The experiment was repeated 3 times and the quantification is shown in the bar graphs above the gels (p<0.001). C. “Hinge’Linker” cohesin trimer ATPase activity measurements. ATP hydrolysis by the “Hinge-Linker” cohesin complex was measured in 4 different conditions. Light shade bars represent WT full length cohesin’s ATPase activity in each condition. D. Sequence alignments of SMC1 and SMC3 showing the conservation of the mutated residues. SMC1 R507,K511,R513,K514 and SMC3 K681,R682,R684,K689 are in black squares. E. In vitro cystein crosslinking to measure the formation of the Zipped up state of WT full length cohesin trimer versus the BDE trimer. The experiment was repeated 3 times and the quantification is shown in the bar graphs above the gels (p<0.001). F. 8DE cohesin trimer ATPase activity measurements. ATP hydrolysis by the truncated cohesin complex was measured in 4 different conditions. Light shade bars represent WT full length cohesin’s ATPase activity in each condition.

We will refer to this region where the hinge domain transitions into the coiled coil as the cohesin ‘wrist’. We noticed that the cohesin wrist region is lined with highly conserved surface residues that form a positively charged patch (Figure 4D). Results described above show that the rigidity of the cohesin wrist region is crucial for zipping-up of the coiled coils at the joint region. We next asked if the conserved positively charged patch in the cohesin wrist is important for the same process. To do this, we chose 8 conserved surface residues (4 on each SMC) that form a positively charged surface patch and engineered a charge reversal mutant, substituting K with D and R with E. We refer to this mutant as 8DE cohesin. We found that this mutant showed a significant defect in the zipping-up of the coiled coils at the joint (Figure 4E) and was capable of hydrolysing ATP in the absence of DNA and to a large extent in the absence of Scc2 (Figure 4F). These results strongly support the notion that the cohesin wrist region is critical for the imposing the constraint on the ATPase domain by causing the zipping-up of the coiled coils at the joint. Moreover, these observations imply that the Scc2 and DNA overcome this constraint to cause unzipping of the coils and thus promote ATP hydrolysis by cohesin.

### Folding of the coiled coils at the elbow does not affect cohesin’s ATPase activity

The experiments described in this study thus far suggest an inverse correlation between the zipping-up of the coiled coils with cohesin’s ATPase activity. Another striking feature of the SMC coiled coils is their folding at the elbow (Burmann et al., 2019). In the absence of ATP, cohesin’s coiled coils while being zipped-up along their length, are also folded at the elbow region (Petela et al., 2021). In the folded conformation, the hinge domain interacts with the joint region of the coiled coils. This interaction has been measured in vivo in yeast cells using BMOE crosslinking of cysteine substituted cohesin (Petela et al., 2021). Most models for LE suppose recurrent rounds of zipping and folding of the coils followed by their unzipping and unfolding during extrusion (Oldenkamp and Rowland, 2022). In order to address if recurrent folding and unfolding of the coiled coils is necessary for LE, a recent study generated an engineered cohesin containing a fusion of the FRB domain into the Smc3 hinge and the FKBP domain to the N-terminus of Scc1 (Bauer et al., 2021). In the presence of Rapamycin this modified cohesin accumulated in a conformation with a sharp bend in the middle of the coiled coils. In this state, the modified cohesin was incapable of hydrolysing ATP in the presence of Scc2 and DNA and consequently incapable of LE (Bauer et al., 2021). However, folding was induced by connecting the hinge domain to the N terminus of the kleisin subunit. There is no evidence for cohesin every existing in such a state. Also, it is unclear what the molecular consequences of artificially tethering the N-terminus of Scc1 to the Smc3 hinge are and whether the Smc3-Scc1 interface, which is critical for cohesin’s ATPase activity, was still intact when folding was induced by addition of rapamycin. Under conditions when cohesin adopts the folded conformation, the hinge domain is in contact with the coiled coils at the joint region and not the N-terminus of Scc1 (Petela et al., 2021). Cohesin has been demonstrated to adopt this conformation in living cells using BMOE mediated crosslinking of cysteine residues place in the hinge and the joint region (Petela et al., 2021). We reasoned that these cysteine substitutions would serve as a better proxy for the natural folding of cohesin’s coils and would serve as a tool to measure the effect of coils folding on the ATPase activity. Therefore, we expressed and purified a form of cohesin that can be crosslinked in its naturally folded conformation (Figure S1A and S1B). Treatment of this version of cohesin with BMOE resulted in about 70% of the molecules being crosslinked in the folded conformation (Figure 5A). Our results with the zipping-up reporter indicate that the formation of the clamped state results in the unzipping of the coils (Figure 1F). We asked if the same would be true for the folding of the coiled coils. Interestingly, we found a significant reduction in the folding of the coils under conditions EQEQ cohesin accumulates in the clamped state (Figure 5B). Moreover, the cohesin wrist mutant that is compromised for the coiled coil zipping-up was also severely defective in the folding of the coils (Figure 5C). These results are consistent with previous observations with single molecule FRET (Bauer et al., 2021) and suggest that coordinated folding and zipping-up of the coiled coils inhibit cohesin’s ATPase. They also raise an attractive possibility that the folding of the coils stabilises the zipped-up conformation or vice versa. A clear prediction therefore would be that folding of the coils would impede ATP hydrolysis just like the zipping-up of the coils at the joint does. To test if this is the case, we measured ATP hydrolysis by cohesin that was pre-crosslinked in folded conformation (Figure 5D). As observed before, ATP hydrolysis by the wild type cohesin that does not contain any cysteine substitutions was unaffected by BMOE treatment. Cohesin with the fold cysteines was efficiently crosslinked by BMOE, with 59% of the cohesin molecules crosslinked in the folded conformation. Incredibly, we found that irreversible folding of about two thirds of cohesin molecules did not cause any measurable change in ATP hydrolysis. This striking result suggests that the folding of the coiled coils at the elbow does not impede cohesin’s Scc2 and DNA dependent ATPase activity. Clearly, cohesin can hydrolyse ATP continuously without having to go through repeated cycles of folding and unfolding of the coiled coils. This raises an interesting question, are the cohesin complexes trapped in the folded state capable of LE?

**FIGURE 5.**
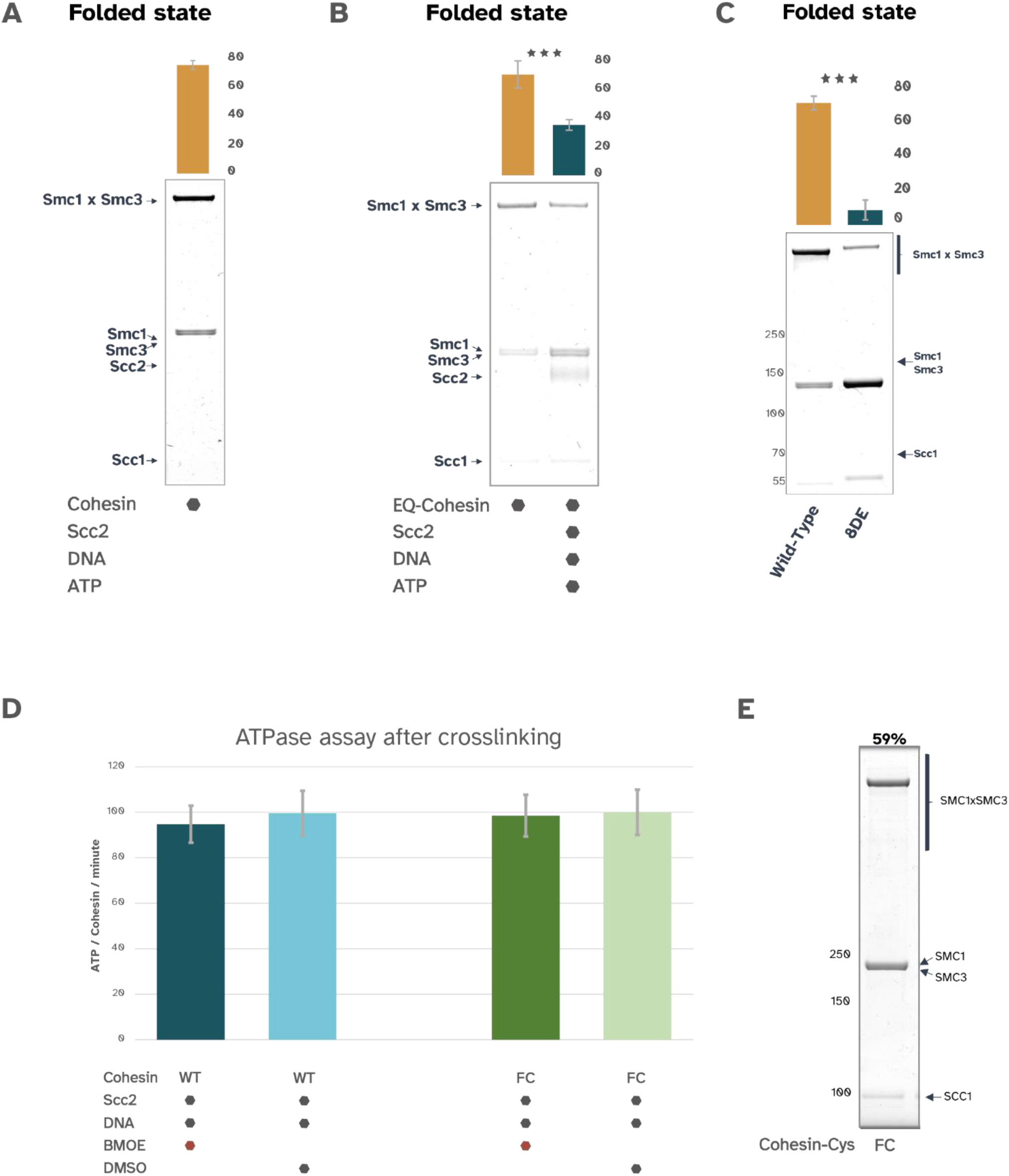
A. In vitro cystein crosslinking of Smc1R578C and Smc3V933C measuring the Smc1 hinge - Smc3 ascending coil association indicative of the folded confromation. B. In vitro cystein crosslinking to measure the formation of the Folded state of EQ cohesin trimer in the presence or absence of Scc2, DNA and ATP. The experiment was repeated 3 times and the quantification is shown in the bar graphs above the gels (p<0.001). C. In vitro cystein crosslinking to measure the formation of the Folded state of WT full length cohesin trimer versus the 8DE trimer. The experiment was repeated 3 times and the quantification is shown in the bar graphs above the gels (p<0.001). D. ATPase activity measurements after crosslinking with BMOE in comparison with DMSO controls. Cohesin trimers with cysteine pairs capturing the folded state (UC-elbow) and were tested along with (WT). Each experiment was repeated 3 times (p<0.001). E. n vitro cystein crosslinking to measure the efficiency of the crosslinking in the folded state (FC) before release in the ATPase assay.

In summary, our in vitro analysis of cohesin has provided us key insights into the relationship between cohesin’s ATPase cycle and the conformation of the SMC coiled coils. In the absence of ATP, cohesin’s coiled coils are zipped-up along their length and folded at the elbow (Figure 5D). While ATP is sufficient to induce engagement of the ATPase domains, the bound ATP molecule cannot be hydrolysed without Scc2 and DNA. Presumably, in the absence of ATP, the ATPase heads can go through cycles of engagement and disengagement without ATP hydrolysis (Bauer et al., 2021) (Figure 5D). Zipping-up of the coiled coils at the joint region imposes a constraint on the ATPase heads and impedes hydrolysis. Clamping of DNA over the ATPase heads by Scc2 leads to unzipping of the coiled coils, at least up to the joint region. This change in conformation is necessary for ATP hydrolysis (Figure 5D). Most models for LE involve recurrent cycles of folding and unfolding coordinated with zipping-up and unzipping of the coiled coils. Interestingly, our data imply that neither the permanent zipping-up of the coils at the elbow nor indeed the folding of the coils at the elbow impede ATP hydrolysis. This raises the question as to whether such conformation changes are necessary for LE.

## Materials and methods

### Reagents

Reagents:
|  |  |  |  |
| --- | --- | --- | --- |
| Chemical compound, drug | ATP Lithium Salt | Sigma | Cat# 11140965001 |
| Chemical compound, drug | Bismaleimidoethane (BMOE) | ThermoFisher | Cat# 22323 |
| Chemical compound, drug | Complete EDTA free protease inhibitor cocktail | Roche | Cat# 4693132001 |
| Chemical compound, drug | Cre Recombinase | New England Biolabs | Cat# M0298S |
| Chemical compound, drug | Desthiobiotin | Fisher Scientific | Cat# 12753064 |
| Chemical compound, drug | DMSO | Merck Life Science UK Limited (Sigma) | Cat# D8418-50ML |
| Chemical compound, drug | DTT | ThermoFisher | Cat#R0861 |
| Chemical<br>compound,<br>drug | EtBr | ThermoFisher | Cat#15585011 |
| Chemical<br>compound,<br>drug | FuGENE HD<br>Transfection reagent | Promega | Cat# E2311 |
| Chemical<br>compound,<br>drug | Gibson Assembly Mix | New England<br>Biolabs | Cat# E2611L |
| Chemical<br>compound,<br>drug | NuPAGE 3–8% Tris-Acetate Protein Gels | ThermoFisher | Cat# EA0378BOX |
| Chemical<br>compound,<br>drug | PMSF | Sigma | Cat# 329-98-6 |
| Chemical<br>compound,<br>drug | Quick Coomassie Stain | Generon | Cat# GEN-QC-STAIN |
| Chemical<br>compound,<br>drug | Sf900 II SFM | ThermoFisher | <u>Cat# 10902104</u> |
| Chemical<br>compound,<br>drug | Supernuclease | SinoBiological | Cat# SSNP01 |
| Chemical<br>compound,<br>drug | TCEP | ThermoFisher | Cat# 20490 |
| Chemical compound, drug | 4xLDS | ThermoFisher | Cat# NP0007 |
| Commercial assay or kit | HiLoad 16/60 Superdex 200 | GE Healthcare | Cat# GE28-9893-35 |
| Commercial assay or kit | HiTrap Q HP | GE Healthcare | Cat# GE29-0513-25 |
| Commercial assay or kit | StrepTrap HP | Fisher Scientific | Cat# 11540654 |
| Commercial assay or kit | Superose 6 Increase 10/300 GL | VWR | Cat# 29-0915-96 |
| Commercial assay or kit | EnzChek phosphate assay kit | Invitrogen | Cat# E6646 |

### Plasmids

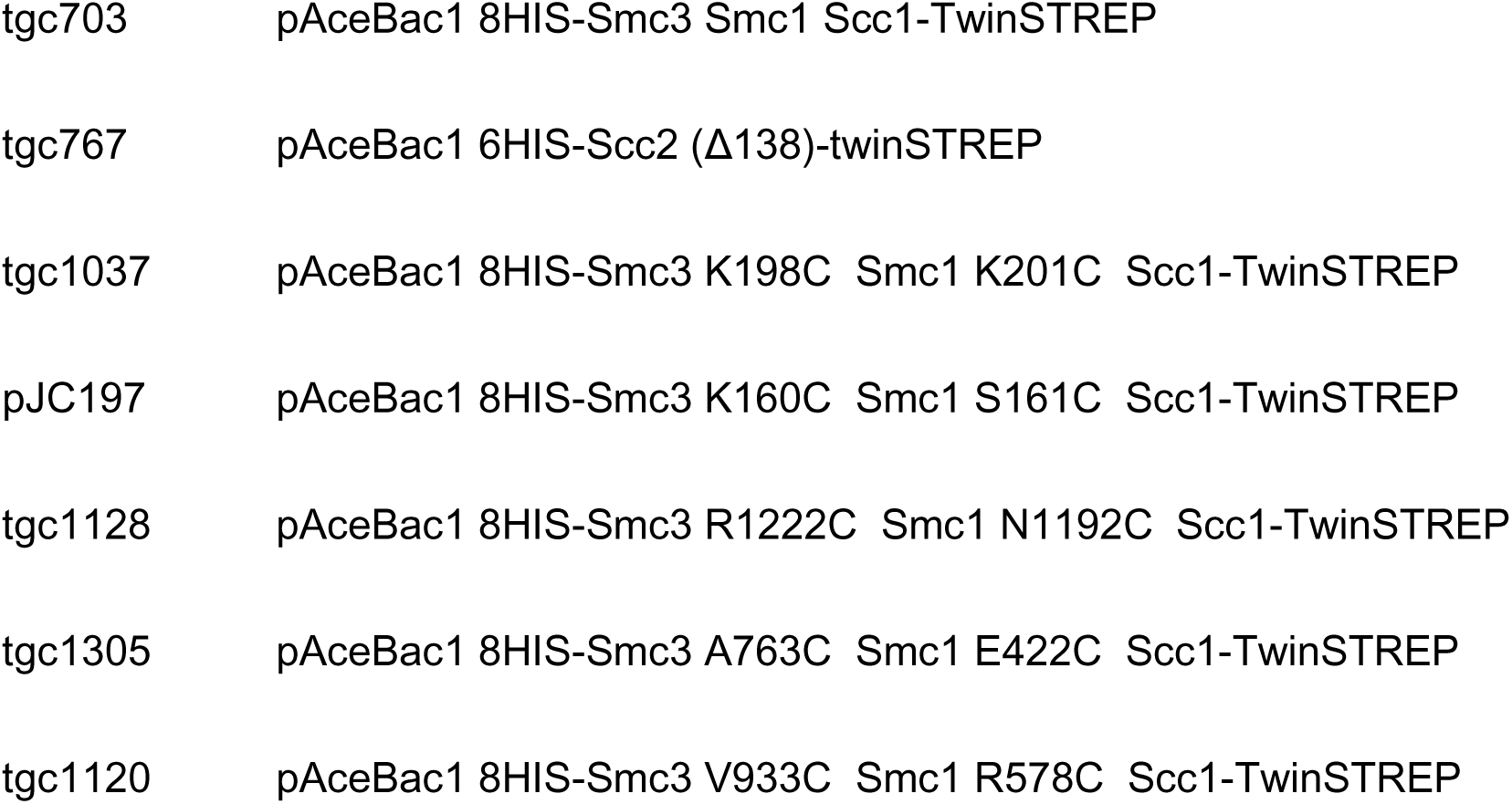
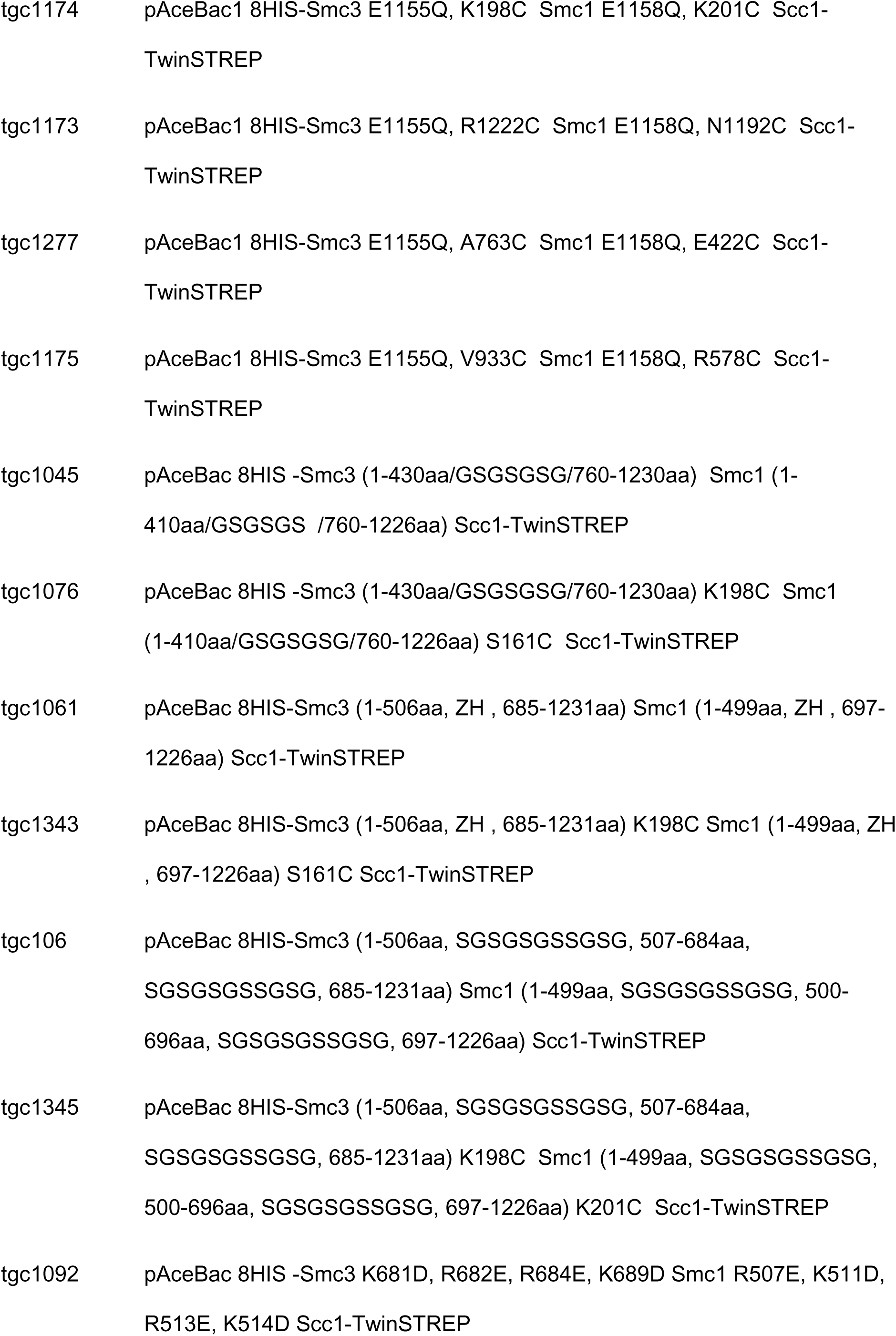
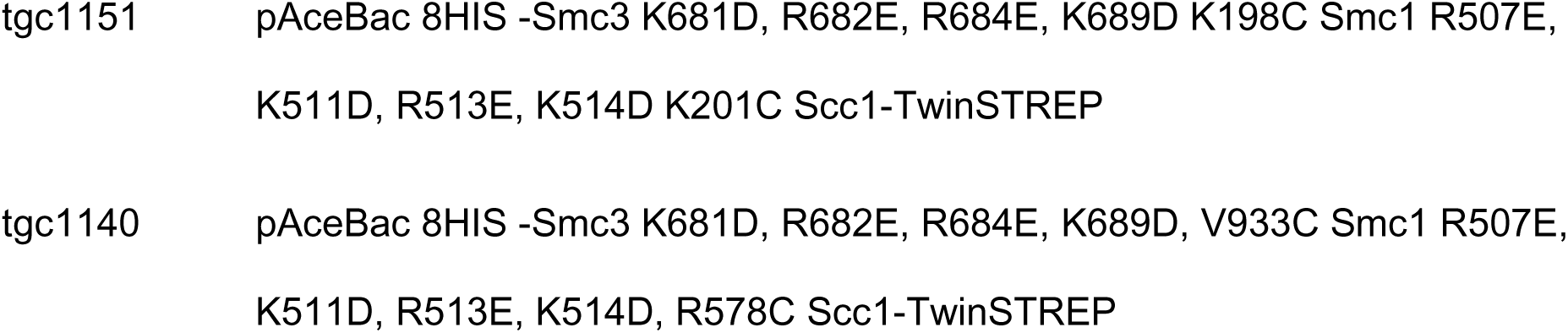

### Recombinant yeast cohesin complex cloning

The S. cerevisiae cohesin genes were codon optimized for SF9 cell expression and synthesized using Thermo Fisher’s Genescript service. Vectors from the multibac expression protocol were used for cloning. Smc1, 8His-Smc3, 6His-Scc2(139-1493)-2xStrepII, 2xStrepII-Scc3 and variations thereof were cloned in the pAceBac vector, while Scc1-2xStrepII was cloned in the piDC vector. pAceBac plasmids were amplified in Top10 cells while piDC plasmids required Pir+ cells. 8His-Smc3 and Smc1 were both inserted in one pAceBac vector and expressed together. Any truncations, mutations or domain insertions for constructs were done using the Gibson assembly protocol. A GS linker was used for the truncated complexes. Plasmids containing all 3 cohesin trimer proteins were made by cre recombination of the pAceBac plasmid containing both SMCs and the piDC containing Scc1.

### Baculovirus generation

Vectors containing the genes of interest were transformed into DH10 cells where transposition into the bacmid was confirmed by blue white screening and antibiotic selection. Following lysis of the cells, isopropanol and ethanol precipitation was used for the isolation of the bacmid at >2μg/ml concentration. 2μg of bacmid were transfected to 2ml of SF9 cells at 0.5 x 106/ ml concentration using the FugeneHD reagent. The cells were grown in SF900II SFM at 27oC in 6 well plates for 3 days. Upon visual examination and confirmation of viral infection and ceasing of cell growth, the cells were span at 2000g. The P1 virus was collected as the supernatant. Amplification for P2 virus was achieved by infecting 50ml of SF9 cells at 2×106/ml concentration with 500μl of P1. P2 was collected after 3 days upon confirmation of infection.

### Protein Expression

For the expression of the protein of interest, 500ml of SF9 cells at 2×106/ml concentration were infected with 5ml of P2 virus. The cells were incubated at 27oC in suspension shaking at 120rpm for 2 days. After 2 days the culture was sampled and evaluated for viability and expression levels. Upon confirmation of protein expression the pellet was collected by centrifugation at 2000g for 15min washed ones with PBS buffer and then flash frozen with liquid nitrogen. The pellet was stored at −80oC.

### Protein purification

Cell pellets were thawed at 4oC and suspended in 100ml buffer A (50mM Hepes pH 7.5, 150mM NaCl, 1mM TCEP, 5% Glycerol) supplemented with 1 complete EDTA free Protease inhibitor tablet, and 100units of Supernuclease. Lysis was achieved via sonication using the Vibra Cell Sonicator (VCX130PB-1) for 2 minutes (10 sec intervals) at 80% amplitude. 1mM PMSF was added after sonication and the lysed cells were incubated for 15min on ice. Finally the solution was centrifuged at 50.000g in 4oC to pellet any cell debris while the supernatant was collected and passed through a sterile 0.8μM pore filter. The protein of interest was then purified following a 3-step protocol, including Affinity chromatography, Anion exchange chromatography and Size exclusion chromatography. Affinity pulldown was performed using the 5ml Strep-Trap HP column, connected to an Akta Explorer apparatus. The column was washed with Buffer A and eluted with 15ml buffer A supplemented with 5mM d-desthiobiotin. The elution was then diluted to 30ml using Buffer B (50mM Hepes pH 7.5, 100mM NaCl, 1mM TCEP, 5% Glycerol) and was then loaded to the HiTrap Q HP anion exchange column. Elution was done across a gradient of 100-1000mM NaCl. Finally size exclusion chromatography was used to select against any aggregates and the protein was collected and flash frozen in liquid nitrogen. In the case of the isolated hinge constructs, Hi-Trap Talon column was used to pull down on the 8His tag on Smc3 and the ion exchange step was omitted due to the high isoelectric point.

### DNA oligo duplex

dsDNA was created by annealing a pair of complementary 40bp oligos (GAATTCGGTGCGCATAATGTATATTATGTTAAATAAGCTT, AAGCTTATTT AACATAATATACATTATGCGCACCGAATTC). The 100mM oligos were mixed at 45μl each, with 10μl NEB buffer 2. The annealing reaction was done in a thermocycler, first heating the sample at 95oC, and then gradually decreasing the temperature to 10oC in 0.1oC increments. The final concentration of the dsDNA was 45mM.

### ATPase assay

ATPase activity was measured using the Enz-Chek phosphate assay kit as described in (Voulgaris and Gligoris, 2019).

### Protein Crosslinking Assay

Cohesin trimers with Cysteine pairs were incubated at 150nM concentration with oligo duplex DNA at 450nM, ATP at 10nM and Scc2 at 150nM. The reactions were incubated for 5 minutes in room temperature. BMOE was then added to a final concentration of 1mM and the reaction was incubated for 5 minutes on ice. The sample was then denaturated with 4xLDS buffer and boiled at 95oC for 5 minutes. The results were visualized at 3-8% Tris-Acetate gels stained with Quick Coomassie stain.

### Sequence alignment

Protein sequences were aligned using the Clustal Omega tool in the EMBL-EBI website and visualized using the Jalview software.

---

**FIGURE S1.**
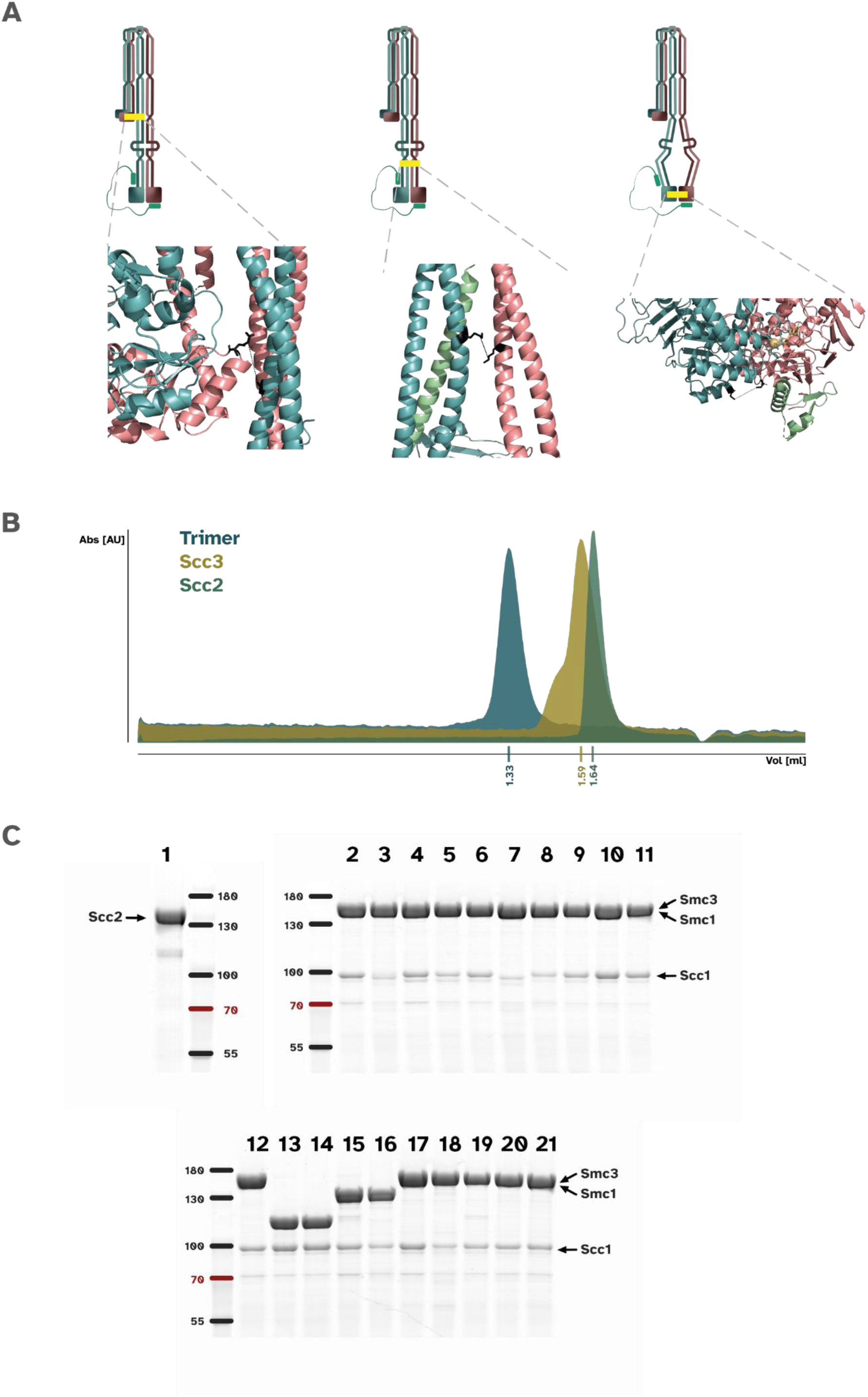
A.
i. Schematic showing the crosslinkning pair (yellow bar) to detect the folded state of cohesin. The positions of the Cysteins [Smc1 R578C Smc3 V933C]are shown in the blown up structure in black.
ii. Schematic showing the crosslinkning pair (yellow bar) to detect the zipped up state of cohesin.The positions of the Cysteins [Smc1 K201C Smc3 K198C]are shown in the blown up structure in black.
iii. Schematic showing the crosslinkning pair (yellow bar) to detect the head-engaged state of cohesin.The positions of the Cysteins [Smc1 N1192C Smc3 R1222C]are shown in the blown up structure in black B. Size exclusion chromatography profile of purified cohesin trimer (blue), Scc3, (yellow) and Scc2 /l 150 (green) C. Coommassie stained SDS-page gel of all the proteins used in the study.
1. Scc2
2. WTTrimer
3. WT Trimer/ Zipped-up pair cysterines
4. WT Trimer/ J-heads pair cysterines
5. WT Trimer/ E-heads pair cysterines
6. WT Trimer *I* Upper-coil pair cysterines
7. WT Trimer *I* Folded pair cysterines
8. EQ Trimer/ Zipped-up pair cysterines
9. EQ Trimer/ E-heads pair cysterines
10. EQ Trimer/ Upper-coil pair cysterines
11. EQ Trimer/ Folded pair cysterines
12. WTTrimer
13. Truncated Trimer (elbow)
14. Truncated trimer *I* Zipped-up pair cysteines
15. “Zink-hook” trimer
16. “Zink-hook” trimer *I* Zipped-up pair cysteines
17. “Hinge-linker’’ trimer
18. “Hinge-linker’’ trimer/ Zipped-up pair cysteines
19. BDE Trimer
20. BDE Trimer/ Zipped-up pair cysteines
21. BDE Trimer/ Folded pair cysteines Ladder [NEB PageRuler]

